# Limitations of compensatory plasticity: the organization of the primary sensorimotor cortex in foot-using bilateral upper limb dysplasics

**DOI:** 10.1101/190462

**Authors:** Ella Striem-Amit, Gilles Vannuscorps, Alfonso Caramazza

**Author notes:** Corresponding author: Ella Striem-Amit, Department of Psychology, Harvard University, Cambridge, MA 02138 USA,.

## Abstract

What forces direct brain organization and its plasticity? When a brain region is deprived of its input would this region reorganize based on compensation for the disability and experience, or would strong limitations of brain structure limit its plasticity? People born without hands activate their sensorimotor hand region while moving body parts used to compensate for this ability (e.g. their feet). This has been taken to suggest a neural organization based on functions, such as performing manual-like dexterous actions, rather than on body parts. Here we test the selectivity for functionally-compensatory body parts in the sensorimotor cortex of people born without hands. Despite clear compensatory foot use, the sensorimotor hand area in the dysplasic subjects showed preference for body parts whose cortical territory is close to the hand area, but which are not compensatorily used as effectors. This suggests that function-based organization, originally proposed for congenital blindness and deafness, does not apply to cases of the primary sensorimotor cortex deprivation in dysplasia. This is consistent with the idea that experience-independent functional specialization occurs at relatively high levels of representation. Indeed, increased and selective foot movement preference in the dysplasics was found in the association cortex, in the inferior parietal lobule. Furthermore, it stresses the roles of neuroanatomical constraints such as topographical proximity and connectivity in determining the functional development of brain regions. These findings reveal limitations to brain plasticity and to the role of experience in shaping the functional organization of the brain.

**Significance Statement:** What determines the role of brain regions, and their plasticity when typical inputs or experience is not provided? To what extent can extreme compensatory use affect brain organization? We tested the functional reorganization of the primary sensorimotor cortex hand area in people born without hands, who use their feet for every-day tasks. We found that it is preferentially activated by close-by body-parts which cannot serve as effectors, and not by the feet. In contrast, foot-selective compensatory plasticity was found in the association cortex, in an area involved in tool use. This shows limitations of compensatory plasticity and experience in modifying brain organization of early topographical cortex, as compared to association cortices where function-based organization is the driving factor.

**Classification:** Biological Sciences\Neuroscience

## Introduction

What determines the role of brain regions, and their plasticity when typical inputs or experience is not provided? To what extent can extreme compensatory use affect brain organization? Or is brain organization limited in its plasticity, due to strong neuroanatomical constraints? The balance between nature and nurture has been a long-standing question in neuroscience.

Here we tackle this question using the model of limb absence, in the sensorimotor cortex of upper limb dysplasic individuals, born without hands. What would take the place of the missing hands? Several recent studies stress the role of experience-based plasticity, and claim that this region reorganizes to support the use of other body parts to perform everyday actions. People born without hands showed activation of the hand sensorimotor cortex for body parts used to “replace” congenitally missing hands, be it the feet (1) or other, multiple body parts (in addition to the remaining hand) in congenital one-handers (2). In the somatosensory cortex, the hand responsive area shrinks depending on the size and use of the hand remains (3, 4), and foot somatosensory stimulation activated the lateral sensory cortex (5). Furthermore, in two individuals born without hands bilaterally, a study has shown that strong stimulation of the lateral motor cortex generated motor evoked potentials not only in the residual finger or shoulder of the subjects but also in the foot, and interfered with performing a foot motor task (1). These findings were interpreted as evidence for robust plasticity of the sensorimotor cortex and functional takeover of the hand area by the feet. Furthermore, they raised the possibility that the primary sensorimotor cortex is functionally-selective rather than selective for topographical body parts, such that the hand area may correspond to any effector that functions as a hand in everyday tasks like grasping and manipulating objects (2).

However, the *specificity* of such supposed compensatory reorganization (for the body part now used as hands) has not been thoroughly tested. None of the studies tested and showed that the hand area is more activated for compensatorily used body parts than for other proximal, but non-compensatorily used organs. Is the hand sensorimotor cortex indeed selective for compensatorily used body parts which serve as effectors? Alternatively, plasticity due to compensatory effector experience may be limited by neuroanatomical constraints such as topographical proximity and connectivity of this brain system, enabling only a takeover by closer cortical territories or contralateral intact body parts, akin to what is found in late-onset amputation (6-11).

Here we test these competing hypotheses by mapping sensorimotor responses to movement of various body parts in five individuals born without hands (dysplasic subjects), who use their feet for everyday functions. The implications of the results are discussed in the context of the broader issue of retained functional specialization in congenital blindness and deafness.

## Results

To test the specificity of plasticity in the sensorimotor hand area in people born without arms or hands (see **Table 1**), we used an active motor paradigm, designed to activate both primary somatosensory and primary motor cortices, similar to previous studies of reorganization (2, 12-15). We scanned five dysplasic subjects, as well as a control group, as they performed simple flexing movements of different parts of their body. The body parts chosen for this experiment included not only the hands (in the controls, to be used as a localizer), but also the feet, which the dysplasic subjects use to overcome their disability. Our study participants, according to self-report, rely largely on their dexterous feet (dominantly their right foot; all were right footed) to perform daily typical manual activities. Their feet are extraordinarily dexterous, allowing them to use cell phones, utensils and nearly all other everyday tools (see list at **Table S1**). In a questionnaire of tool use, all the dysplasic subjects reported to use the clear majority of the tools they have used with their lower limbs (see **Fig. S1**). Foot tool-use accounted for a minimum of 92% of the used tools, although a minority of tools were jointly manipulated by the lower face or remaining upper limbs in specific individuals (see **Fig. S1**, **Table 1**). Additionally, we inspected movement of the shoulder and lips, expected to activate neighboring cortical regions on both sides of the missing hand territory, as well as a control body part, which is not an immediate cortical neighbor of the hand and is not used compensatorily to replace hand function as a dexterous effector: the abdomen. While abdominal core muscles may be used to stabilize the body and may thus be used in excess by the dysplasics while using their feet, they cannot be used to replace hand function.

**Table 1:**
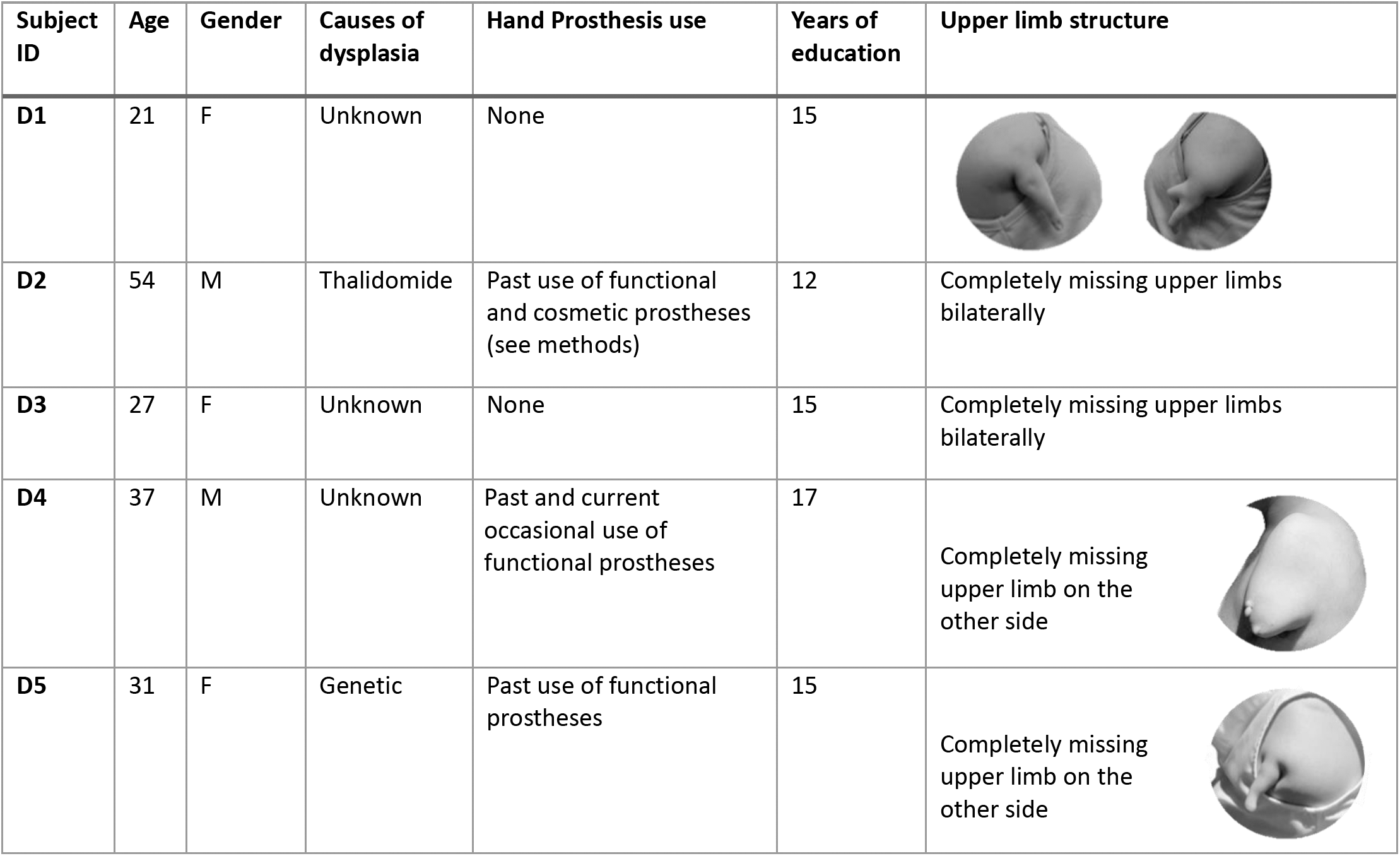
Characteristics of the dysplasic subjects

| Subject ID | Age | Gender | Causes of dysplasia | Hand Prosthesis use | Years of education | Upper limb structure |
| --- | --- | --- | --- | --- | --- | --- |
| D1 | 21 | F | Unknown | None | 15 |  |
| D2 | 54 | M | Thalidomide | Past use of functional and cosmetic prostheses (see methods) | 12 | Completely missing upper limbs bilaterally |
| D3 | 27 | F | Unknown | None | 15 | Completely missing upper limbs bilaterally |
| D4 | 37 | M | Unknown | Past and current occasional use of functional prostheses | 17 | Completely missing upper limb on the other side |
| D5 | 31 | F | Genetic | Past use of functional prostheses | 15 | Completely missing upper limb on the other side |

In the control group, this protocol resulted in a typical somatotopic activation pattern with sensorimotor responses along a superior-inferior axis for the foot, abdomen, shoulder, hand and lips movements (**Fig. 1A**; hand peak is delineated in white), replicating the known Penfield homunculus (16). In the dysplasic subjects, though the peak responses remained topographic, movement in every tested body part extended towards the deprived area and generated some activation in the hand region (**Fig. 1B**). This included also increased activation of the hand area while they moved their right foot, used by these subjects to perform typically manual actions, as reported before. Indeed, plotting the group differences showed that the sensorimotor hand area had stronger activation in the dysplasic subjects than in the controls for moving the various body parts, including the foot (**Fig. 1C,D**).

**Figure 1:**
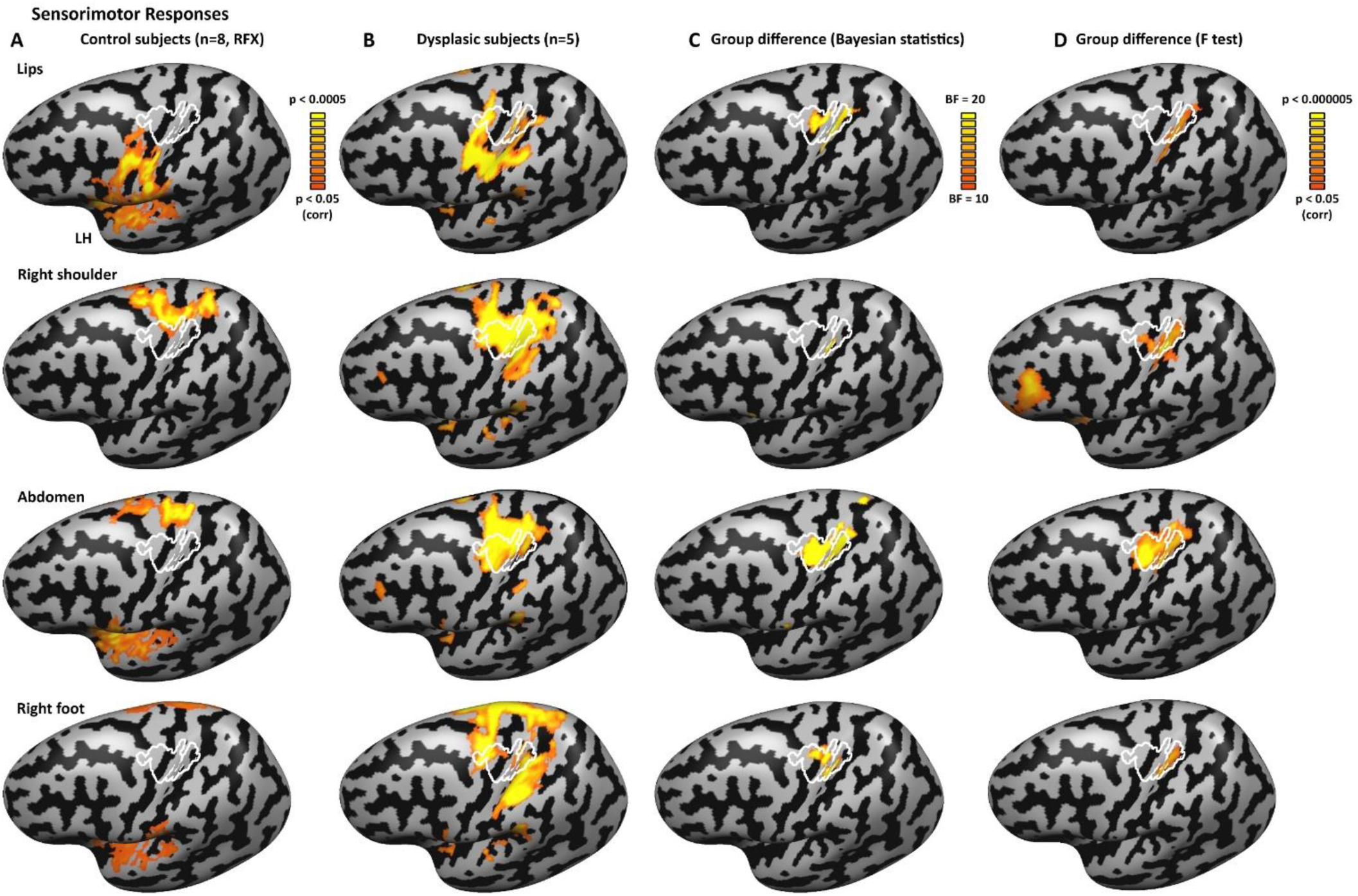
Activation for multiple body parts in the sensorimotor hand area in dysplasic subjects (born without hands) **A.** The activation for body part movement (contraction of the lips, right shoulder, abdomen and right foot) in the typically developed control subject group (random effect GLM analysis; p < 0.05 corrected for multiple comparisons) is shown on the left cortical hemisphere, following the standard Penfield homunculus. The sensorimotor hand area, delineated in white, represents the core area activated by right hand movement in all the control participants (each at p < 0.05 corrected), to account for intersubject variability. **B.** Activation for body part movement is shown for the dysplasic individuals, born without hands (fixed effect GLM analysis, p < 0.05 corrected; see **Table 1** for subject upper limb structure). Movement of each of the tested body parts elicited activation in the hand area to some extent, including, as previously reported, movement of the right foot. **C-D.** Group difference in activation of body part movement is shown in both Bayesian analysis (panel **C**; Bayes factor, BF_10_ > 10 represents strong evidence for the existence of a group difference) and frequentist statistics (panel **D**; mixed effects F test). The sensorimotor hand area is activated more by the dysplasic group for multiple body parts. Group differences extend somewhat more inferiorly in the posterior somatosensory cortex (postcentral gyrus) as compared to the motor cortex, in accordance with a larger cortical representation to the upper limbs in this area (55).

When exploring selectivity of sensorimotor responses, plotting the preferential activation per cortical vertex (in a winner-takes-all approach; **Fig. 2A, B**), the dysplasic subjects show a preference for shoulder and abdomen movements in the typical hand area (**Fig. 2B**; hand area delineated white), which seem to have been displaced and expanded towards the hand area as compared to the activation pattern in the controls. These patterns were consistent across the individual subjects (**Fig. S2**) and found also for the right hemisphere for movement of the contralateral body parts (see **Fig. S3** for comparable analyses of the right hemisphere). We further sampled the response pattern within the hand region-of-interest (ROI; defined by significant overlapping activation for hand movement as compared to baseline in all control subjects). The activation in the dysplasics hand region is strongest for the movement of the shoulder, which is proximal to the missing hand (and therefore also activates the hand region in controls; see also **Fig. 1A, Fig. 2C**), but is also significant for the abdomen, whereas moving the foot does not generate strong significant activation in the hand area at large (**Fig. 2C**). Despite the preferential compensatory use of the right foot in these subjects to overcome their disability, movements of the foot do not seem to favorably overtake this region. In fact, movement of the shoulder and abdomen activates the hand region in the dysplasics significantly more than foot movement (p < 0.005 for both). Similarly, in a whole brain analysis, when contrasting moving the foot as compared to the non-compensatory but more proximal abdomen, the dysplasic subjects do not show significant activation in their sensorimotor hand area, and activity was found for the reverse contrast (**Fig. 2D**; see **Fig. S5A,C** for unsmoothed individual subjects data and overlap analysis).

**Figure 2:**
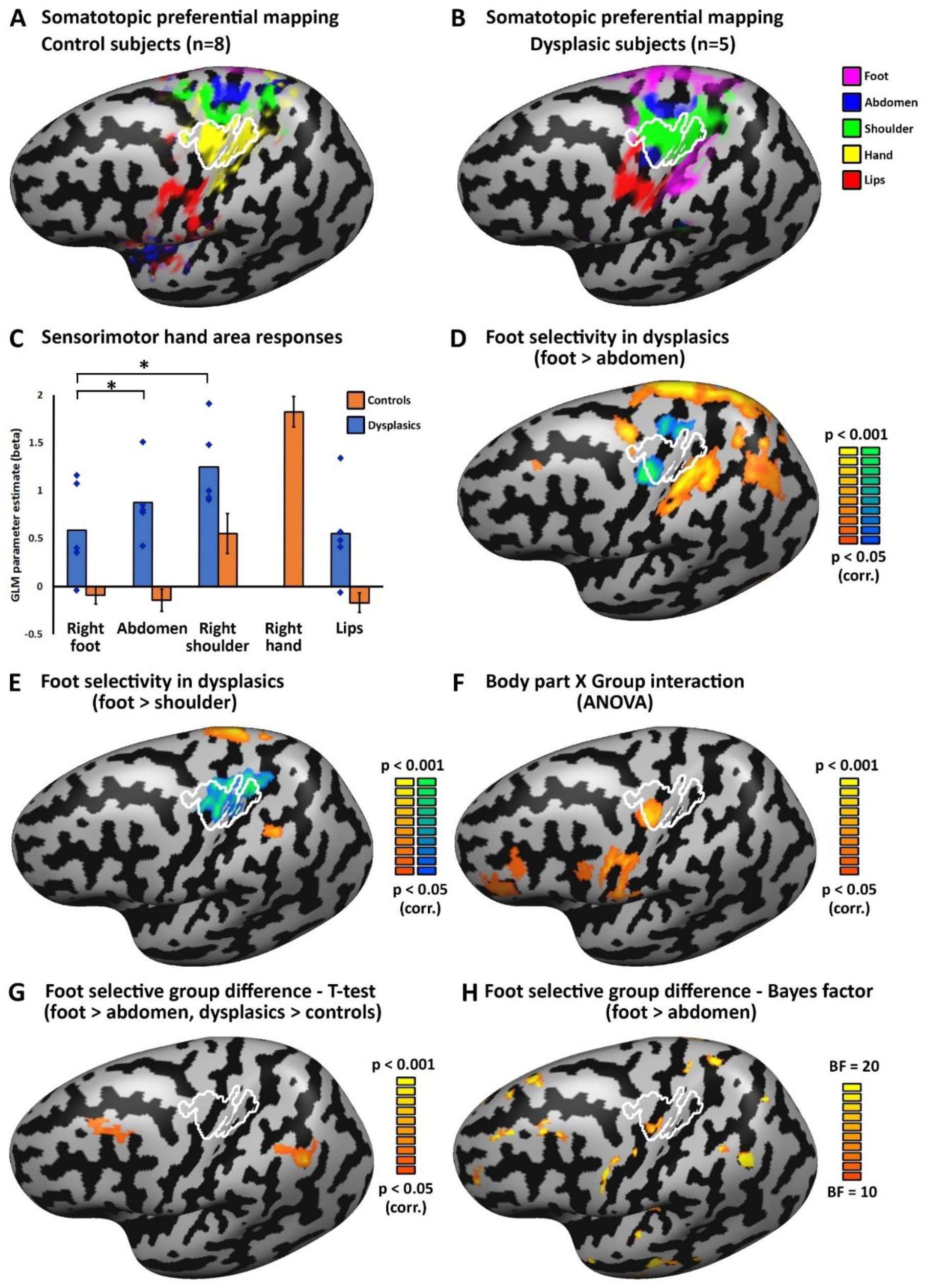
The sensorimotor hand area in the dysplasics does not show selectivity for the compensatorily used foot. **A.** Preferred body part responses for contraction movements (winner-takes-all approach) for the control subjects follows the standard Penfield homunculus. The sensorimotor hand area is delineated in white, representing the core area activated by right hand movement in all the control participants (each at p < 0.05 corrected), to account for inter-subject variability. **B.** Preferred body part responses for contraction movements for the dysplasic group shows a preference for shoulder (and to some extent abdomen, in the motor cortex) movements in the hand area, despite the extensive use of the feet to perform typically manual fine-motor tasks. For individual subject maps showing high reproducibility of this effect see **Fig. S2**. The same is found in the right hemispheres for movement in the contralateral side of the body: see **Fig. S3**, and in passive tactile stimulation of the body; see **Fig. S4**). Curiously, preferential activation for abdomen movement was found also on the anterior inferior border of the hand region, in agreement with evidence of a potential discontinuity in the motor cortex surrounding the hand area (56, 57). **C.** Sensorimotor responses were sampled from the hand area, showing that this region in the dysplasics is more activated by proximal body parts (shoulder and abdomen/trunk) than by foot movements (p < 0.005 for both comparisons). Error bars for the control group (orange bars) represent standard error of the mean. Individual data points (blue diamonds) are presented for the five dysplasic individuals in addition to the group average. **D.** Foot movement selectivity (over abdomen movement, representing a control body part that does not serve compensatorily as an effector) in the dysplasics can be found in the superior parietal lobule and premotor cortex, but not in the hand primary sensorimotor cortex, which shows the reverse preference. For individual subject maps see **Fig. S5**. **E.** Movement selectivity comparing the shoulder and foot in the dysplasics shows a robust preference to shoulder movement (a proximal, non-compensatory body part), rather than to foot movement, in the hand area. For individual subject maps see **Fig. S5**. **F.** Overall body part selectivity (comparing movement of all shared body parts; e.g. lips, shoulder, abdomen and foot) differs between the dysplasics and controls (ANOVA Body part X Group interaction) in the frontal lobe and in the sensorimotor hand area. **G.** A direct comparison of the selectivity to right foot movement (vs. abdomen movement) between the dysplasics and control subjects shows potential for plasticity specific to the compensatorily used foot in the association cortices, in the angular/supramarginal gyri and middle frontal gyrus, but not in the primary sensorimotor cortex. **H.** Bayes factor (BF_10_) for difference between the groups in their differential activation to right foot movement (vs. abdomen movement) is shown. The dysplasics show different selectivity level for right foot movement as compared to the controls in various cortical loci, including the sensorimotor hand area. However, the group difference found in the primary sensorimotor hand area in this analysis reflects a preference in the dysplasics group towards the abdomen movement (compare to panel **D**).

Even more robustly, whole brain analysis contrasting foot movement to shoulder movement shows extensive significant selectivity for shoulder movement in the hand area of the dysplasics (**Fig. 2E**; see **Fig. S5B,D** for individual subjects data and overlap analysis). Preference for the shoulder in the hand area was also found in the somatosensory cortex in a passive somatosensory stimulation experiment (see **Fig. S4** for right and left hemispheres).

This does not mean that no foot selectivity exists in these subjects: when directly contrasting foot movement to movement of the abdomen in the dysplasic subjects (**Fig. 2D**), activation is found not only in the primary sensorimotor cortex foot region which extends laterally from that of the controls, but also in the superior and inferior parietal lobule and posterior superior frontal sulcus. Furthermore, body part selectivity differs between the dysplasics and controls in the frontal lobe and in the sensorimotor hand area itself (ANOVA Body part X Group interaction; **Fig. 2F**). Both frequentist (**Fig. 2G**) and Bayesian statistics (**Fig. 2H**) of foot-selectivity group difference shows additional regions which may be related to compensatory use in the frontal (middle frontal gyrus) and parietal (angular/supramarginal gyrus) lobes. The more sensitive Bayesian analysis hints at group effects also in additional foci and in the temporal lobe, but these may require verification from additional studies, perhaps with increased group size. Therefore, compensatory plasticity related to foot use in the dysplasics is found in the association parietal cortex, in the inferior parietal lobule, and not in the primary sensorimotor hand area.

We then inspected if functional connectivity (FC) may reflect use-based plasticity within the primary sensorimotor cortex. Plotting the FC of the hand area with the cortical areas of the other tested body parts shows that no specific increase in preferential connectivity exists in the dysplasics to the foot region (**Fig. 3A**). The hand-foot FC is not the strongest FC pattern from the hand area, nor is it increased beyond the hand-foot FC of the controls. Furthermore, the group differences in FC from the hand area across the entire cortex shows that the deprived hand area does not seem to change greatly. No group effect was found in an ANOVA mixed effects analysis, and substantial group differences in the FC of the sensorimotor hand area are only evident in the more sensitive Bayesian analysis in the inferior parietal lobule (**Fig. 3B**). No group difference was found in the hand bilateral FC pattern (between the right and left hand ROIs) in the dysplasics, as may be expected due to the bilateral deprivation, nor did they show increased FC between the hand and global signal. Overall, our data does not indicate large-scale changes in FC due to full bilateral congenital upper limb dysplasia.

**Figure 3:**
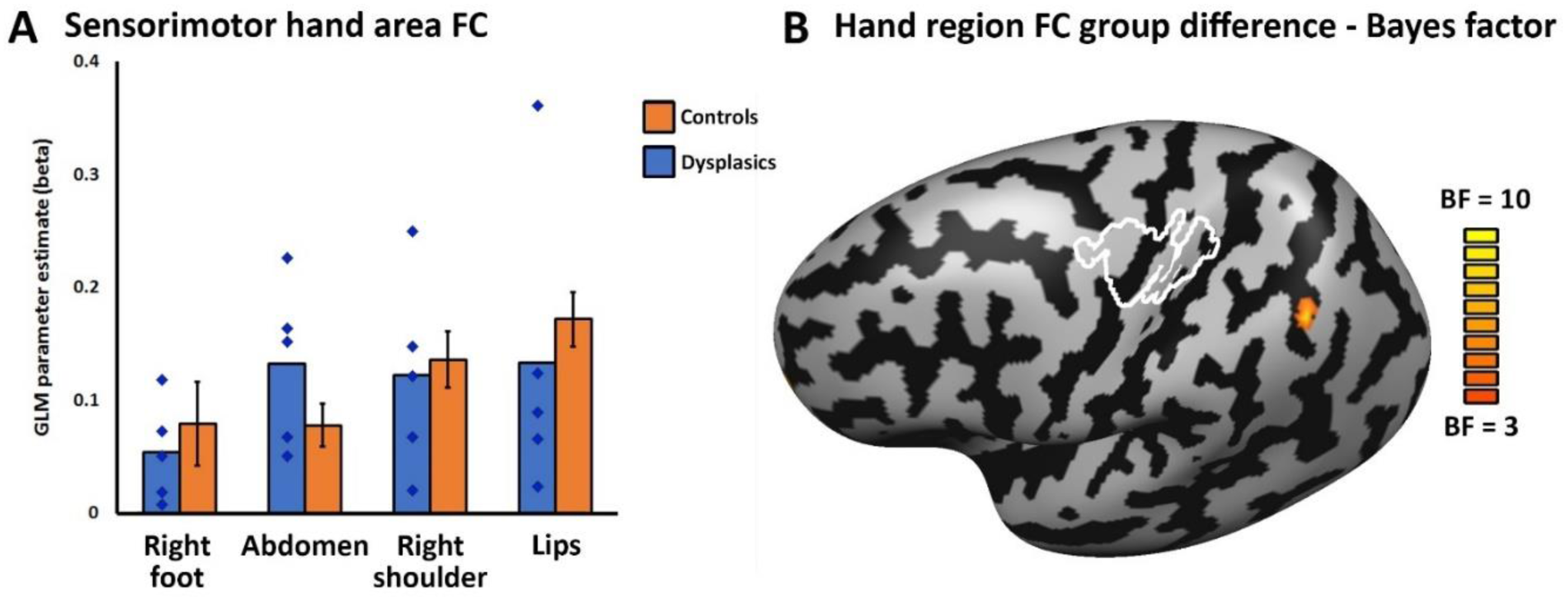
The sensorimotor hand area in the dysplasics does not show enhanced functional connectivity to the compensatorily used foot region. **A.** Functional connectivity between the sensorimotor hand area and sensorimotor areas for the right foot, abdomen, right shoulder and mouth (defined per group) are shown for each group. The hand sensorimotor cortex of the dysplasics does not show increased functional connectivity to the foot area. Error bars for the control group represent standard error of the mean. Individual data points are presented for the five dysplasic individuals. **B.** Bayes factor (BF_10_) for difference between the groups in their functional connectivity from the sensorimotor hand area is shown, not revealing any increased connectivity to foot sensorimotor areas, or otherwise any strong connectivity differences between the groups, except at the inferior parietal lobule. No group effect found in an ANOVA mixed effects analysis.

## Discussion

We tested people born without hands who use their feet to perform every-day manual tasks. We found that their sensorimotor hand area is activated by foot movements more than in typically developed control subjects (**Fig. 1**), replicating previous findings of plasticity in congenital limb absence (1, 2, 5). However, in contrast to previous research, we additionally tested for the selectivity of the compensatorily-used effector, and found that the hand area shows no foot selectivity. Instead, the hand area is significantly more activated for proximal but non-compensatory body parts, which are not used as dexterous effectors, such as the shoulder and abdomen (**Fig. 2**).

Past findings of sensorimotor plasticity in people with congenital hand absence were thought to reflect use-dependent, compensatory plasticity for sensorimotor loss (1, 2, 5). These results were interpreted as evidence that the primary sensorimotor cortex shows functional selectivity for performing tasks typically conducted with the hands, and not necessarily for the specific body part (2). This interpretation builds upon a model developed based on association sensory cortex organization in congenital blindness and deafness. In these cases, the roles of association sensory cortex regions appear to be defined not by their commonly-driving sensory modality (“visual cortex”), but by their computational role. This was demonstrated in vision (17-23) for domain selectivity for complex perceptual and functional categories, such as objects, body parts and scenes (23) as well as for functional tasks such as spatial localization (22) and motion perception (24), and has been extended also to functional tasks for audition (25, 26). Such selectivity is retained, albeit via different sensory inputs, even in the absence of original dominant sensory experience. The finding of compensatory reorganization in people born without one hand raised a provocative suggestion that even the early sensorimotor cortex may also be function-specific rather than body-part specific (2). In contrast, our findings of significant preference for body parts not used as effectors instead of compensatory foot selectivity, even in the complete absence of hands and in individuals who show strong compensatory strategies with foot use, point to limitations of the function-specific model with regard to the primary sensorimotor cortex.

The case of the hand primary sensorimotor cortex in dysplasics differs in several respects from the findings in the blind and deaf. Beyond the different sensory modality, these cases represent different stations of the cortical processing hierarchy. Specifically, in the blind and deaf, claims of functionally-selective organization are limited to the associative sensory cortices, whereas plasticity in people born without hands was tested for the primary sensory-motor cortex. Therefore, while it may be that such principles could govern the organization of higher sensorimotor cortices, they do not seem to apply to the first cortical station processing touch and movement, which is governed by topographic mapping of the body. Foot selectivity in the dysplasics, which may be indicative of compensatory plasticity and effector functional-type organization, was found in association parietal cortex. Specifically, areas in the inferior parietal lobule, mainly in the angular/supramarginal gyrus, appear to show selectivity for foot movement as compared to movement of the abdomen and shoulder, more in the dysplasics than in the controls, as well as an increased functional connectivity to the primary sensorimotor hand area (see **Figs. 2B, D, F**, and **Fig. 3B**), suggesting potential compensatory plasticity in this area that plays a role in tool use (27-29) and more broadly in action and function representations (30). Future studies should inspect these areas in relation to relevant tasks, to see if functionally selective organization can be observed there directly.

An additional difference between these cases is the extent of the deprivation of the sensory modality. In complete blindness or deafness there are no competing inputs within the sensory modality into the early sensory cortices, which could take over deprived parts of the topographic organization. Therefore, while the functional role of these regions is still debated (31-33), the topographic organization in the early stations of the hierarchy are retained (34, 35). In higher stations of the cortical processing, the visual and auditory cortices also receive inputs from other modalities and from downstream cortical stations via feedback connectivity, which become more dominant in the complete absence of visual input. These may then drive cortex organization towards similar functions and domains even in the absence of the typical visual features driving this region. However, the case of an absence of one body part leaves intact inputs from proximal body parts in the topographic organization, generating within-modality competition and overtake. Inputs from nearby cortical stations or subcortical nuclei from competing intra-modal inputs which would typically during development encourage differential specialization, sharpening cortical preferences (36). In the deprived cortex, these now drive the developing cortex more strongly than inputs from other functionally significant downstream stations which can direct it towards compensatory roles. The strongest input to the somatosensory hand area territory would naturally come from its topographical neighbors, which are connected through horizontal, direct and indirect (trans-synaptic) connections (37) as well as from close body parts in subcortical nuclei. Similarly, in the motor cortex, overlap of the fields for different body part could enable neighboring body part movement (37). Indeed, the hand area of control subjects is also significantly activated as they move their shoulder (**Fig. 2C**), demonstrating the large extent of activity from the movement and percept of one body part, and the overlap of body part representations in the primary sensorimotor cortex (37, 38). This overlap and activity of the hand area by the shoulder in controls (**Fig. 2C**) may be the cause of the limited reorganization for this body part evident in the relatively small group difference (**Fig. 1C, D;** weaker group difference for the shoulder than for the abdomen). The ability to evoke plasticity and takeover by nearby body parts has been confirmed in multiple studies of the effects of adult-stage amputation or deafferentation (6-9, 38). It follows naturally that the same level of plasticity would be possible in earlier development, and in the case of congenital limb absence, even without being strongly driven by compensatory use.

Importantly, competing inputs from more remote locations which do not share a common function, such as the foot region may have more limited efficient connectivity to the hand area (e.g., despite its proximity, intracortical connectivity typically does not cross the hand-face-border (39-41)). During development, while the somatotopic maps are still being refined and pruned based on activity patterns and use (36, 42), the extent of the maximal horizontal activation is already defined This existing connectivity *proportion*, as compared to that of non-deprived nearby body parts, poses a constraint to potential plasticity and competing inputs over an area even in the case of completely missing body parts and their typical inputs. While some connectivity can be modulated to permit innervation from remote body parts in cases of deafferentation (43), our data suggests it is not sufficient, in congenital full upper limb dysplasia, to evoke preferential takeover. Thus, the current study reveals clear limitations for brain compensatory plasticity and for the role of experience in modifying brain organization.

## Methods

### Participants

Five individuals born with severely shortened or completely absent upper limbs (individuals with upper limb dysplasia; dysplasics 1-5), and eight typically developed control subjects, matched for age (no group difference; p < 0.25) participated in the experiment. The causes of dysplasia were genetic, ototoxic medications (thalidomide) or unknown. See **Table 1** for the summary of the characteristics of the dysplasics, as well as images of their residual limbs. None of the dysplasics had a history of phantom limb sensations or movements, and all were adept at performing everyday actions and tool-use with their feet (see **Table S1** for a list of tools used with the feet). All the dysplasic participants were right-footed, and used their right foot dominantly. Dysplasic Subject D1 had three residual fingers attached to the shoulder (see **Table 1**). Dysplasic Subjects D2 and D3 had bilateral dysplasic malformations with totally missing upper limbs on both sides (a complete absence of arm, forearm, hand and fingers). Dysplasic Subject D4 had a shortened right arm (± 10 cm humerus). Dysplasic Subject D5 had one residual finger attached to the shoulder. The dysplasic individuals D1, D2, D4 and D5, apart from the congenitally missing hands, had a typically developed body. D3 had a shorter right leg (functionally corrected using a below knee leg and foot prosthesis). All participants had no history of psychiatric or neurological disorder, and gave written informed consent in accordance with the institutional review board of Harvard University.

Dysplasic Subjects D1 and D3 report no history of prosthesis use. D2 occasionally used a wood composite prosthesis with locking elbow and hooks controlled by cables attached to leg straps from 3 to 7 years old, a wood composite prosthesis with electronic elbow and three pronged hooks controlled by micro switches in shoulder harness from 7 to 11 years old and a composite prosthesis with myoelectric elbows and cosmetic hands from 11 to 15 years old. D4 used switch-based right and left arms prostheses as a child and still uses occasionally a switch-based right arm prosthesis as an adult. D5 used myoelectric and manual prostheses five hours a day between 3 and 14 years old. All the subjects who have used prostheses report having used these prostheses mainly, if not uniquely, to pull, maintain in place or push objects but not to manipulate, and used objects for their functional use (e.g., eating with a fork) with their feet.

### Experimental design

The motor experiment was carried out in a block design fMRI experiment (see acquisition detail below). Mouth, abdomen and either side hands (for the control subjects), shoulders and feet were moved (simple flexing/contraction movement) in separate blocks (6 s movement and 6 s rest) in randomized order according to an auditory cue (metronome). Flexing of the hands and feet entailed movements of closing of the palm (drawing the fingers together), flexing of the shoulder lifts it slightly, flexing of the abdomen tightens it, and flexing of the lips pursed the lips together. Four flex and relax movements were performed in each block at a frequency of 0.66 Hz. Due to our focus on the compensatory use of the feet, and as all the dysplasic participants were dominantly right-footed, we used the movements of the right hand and foot for further examination, and provide evidence for similar organization of the right hemisphere in response to left hand and foot movement in supplementary **Fig. S3**.

A supplementary somatosensory experiment with four out of the five dysplasic subjects was carried out in a block design fMRI experiment (see acquisition detail below). Lower face (including the lips), either side of the abdomen, shoulders and feet received natural tactile stimulation in separate blocks (6 s touch and 6 s rest) in randomized order. The natural tactile stimulation was preformed manually using a 4-cm-width paramagnetic paint brush to an auditory cue (metronome), by a trained experimenter, as previously used efficiently for somatotopic mapping in typically developed subjects;(44, 45). In each stimulation block, the body surface was stimulated by brushing the subjects’ skin in a back-and-forth movement along the main body axis. Each body part was stimulated 3 times in each of the three runs of the experiment, in randomized order. Eight catch trials in which the brushing direction was perpendicular to the rostral-caudal direction were present in each run of the experiment, requiring response (foot button press; to ensure subjects attention) and were removed from further analysis. Only runs where the subjects responded in 75% of the catch trials were used for analysis, leaving 3 runs for D1, and 2 runs for subjects D2, D4 and D5.

Functional Imaging: The BOLD fMRI measurements were obtained in a Siemens Tim Trio 3-T scanner at the Center for Brain Science at Harvard University and a 6-channel birdcage head coil. Functional images were acquired with a T2*-weighted gradient echo EPI (GE-EPI) sequence that employed multiband RF pulses and Simultaneous Multi-Slice (SMS) acquisition (factor of 3) (46, 47). The SMS-EPI acquisitions used a modified version of the Siemens WIP 770A. We used 69 slices of 2mm thickness. The data in-plane matrix size was 108×108, field of view (FOV) 21.6cm x 21.6cm, time to repetition (TR) = 2000ms, flip angle = 80°and time to echo (TE) = 28ms. The main experiment had three runs of 186 whole-brain images each collected in one functional scan. The first two images of each scan (during the first baseline rest condition) were excluded from the analysis because of non-steady state magnetization.

Separate 3D recordings were used for co-registration and surface reconstruction. 3D anatomical volumes were collected using T1-weighted images using a MPRAGE T1-weighted sequence. Typical parameters were: FOV= 25.6cm X 25.6cm, data matrix: 256×256×256 (1mm iso voxel), TR=2530ms, TE=1.64, 3.5, 5.36, 7.22ms, flip angle = 7°.

Data analysis was performed using the Brain Voyager QX 2.8 software package (Brain Innovation, Maastricht, Netherlands) using standard preprocessing procedures. Functional MRI data preprocessing included head motion correction, slice scan time correction and high-pass filtering (cutoff frequency: 3 cycles/scan) using temporal smoothing in the frequency domain to remove drifts and to improve the signal to noise ratio. No data included in the study showed translational motion exceeding 2 mm in any given axis, or had spike-like motion of more than 1 mm in any direction. Functional and anatomical datasets for each subject were aligned and fit to standardized Talairach space (48).

Anatomical cortical reconstruction procedures included the segmentation of the white matter using a grow-region function embedded in Brain Voyager. The Talairach normalized cortical surface was then used for surface-based alignment conducted across the subjects according to their cortical curvature (sulci and gyri) patterns. All further analyses were conducted in cortical space. Single subject data were spatially smoothed with a two dimetional 4 vertex full-width at half-maximum Gaussian in order to reduce inter-subject anatomical variability for group analysis, whereas unsmoothed data is presented at the single subjects cortical level (**Figs. S2, S5**). Due to the small sample size of the unique dysplasic group, analyses were based on converging evidence from the group analyses (general linear model; GLM; **Fig 1A,B**, **Fig. 2 A,B,D, E** and **Fig. S2A,B,D,E**), single subject level (**Figs. S2, S5**), mixed effects ANOVA (**Fig. 1D**, **Fig. 2.F, G** and **Fig. S2F, H**), and Bayesian analysis (**Fig. 1C**, **Fig. 2H**, **Fig 3B**; see details below for each analysis), to enable an assessment of the consistency of the findings. The minimum significance level of the results presented in this study (both individual and group level analyses) was set at p < 0.05, corrected for multiple comparisons, using the spatial extent method based on the theory of Gaussian random fields, a set-level statistical inference correction (49, 50)).

The hand sensorimotor cortex (delineated in white in **Figs. 1, 2**) was defined according to a full overlap (100%) of activation to right hand flexing across all control subjects (each at p < 0.05 corrected), to overcome potential individual biases. This region was further used to sample movement responses in each group (**Fig. 2C**) and as a seed to compute functional connectivity (**Fig. 2G, H, I**; see detail below). The contrast of foot selectivity vs. abdomen and foot selectivity vs shoulder in the individual dysplasic subjects (**Fig. S3**) are depicted at a threshold of p < 0.05 corrected for multiple comparisons. Somatotopic preferential mapping was computed at a surface level for each dysplasic subject (**Fig. S2**). Each cortical vertex is colored based on the body part whose movement elicited the higherst activation (GLM estimate, beta value).

Group analyses in the control group were conducted in a hierarchical random effects analysis (RFX GLM; (51)) and, in the dysplasics, a fixed-effect GLM was implemented, due to the group size.

For group level somatotopic preferential mapping, group-level activation maps at a surface level were used for each group seperately, and each cortical vertex is colored based on the body part whose movement elicited the higherst average activation across the group (**Fig. 2A, B**). The same analyses were performed for the somatosensory supplemental experiment (**Fig. S4**). Group analyses are presented on the Colin27 brain inflated cortices, to which individual surface (cortical) data was aligned based on the curvature patterns.

Group comparisons were conducted using both frequentist (t-test and mixed effects ANOVA; with Group and Body-part factors; **Fig. 1D**, **Fig. 2F** and **Fig. S2F,H**) and sensitive Bayesian analyses ((52, 53); **Fig. 1C**, **Fig. 2H**, **Fig 3B**), appropriate for testing small samples of unique populations and patients. The Bayes Factor is the probability of the data under one hypothesis relative to the probability of the data given another (H1/H0; H0 signifying no group difference, H1 signifying a difference between the groups) and, therefore, allows evaluating the strength of the evidence in the data for both alternatives. The Bayes factor (BF10) was calculated by first computing a two-samples independent two-tailed t test between the groups on the effect in question (e.g. in **Fig. 1C**, on the activation for each body part movements). BFs were computed based on the resulting t values using the Matlab function t2smpf provided by Sam Schwarzkopf (www.sampendu.wordpress.com/bayes-factors; (53)). The JZS prior was selected (53), with the default Cauchy prior width r = 0.707. Bayes factor of over 3 is considered substantial evidence and BF over 10 is considered strong evidence against the null hypothesis (52), in our case suggesting a group difference.

### Functional connectivity data analysis and MRI acquisition

A dataset of spontaneous BOLD fluctuations for the investigation of intrinsic (rest state; (54)) functional connectivity was collected while the subjects lay supine in the scanner without any external stimulation or task. The pulse sequence used was gradient-echo EPI with parallel imaging (factor of 4). The data in-plane matrix size was 108×108, field of view (FOV) 21.6cm x 21.6cm, time to repetition (TR) = 1500ms, flip angle = 75°and time to echo (TE) = 28ms. 68 slices of 2mm thickness (with 0.2mm spacing) were used to obtain full coverage of the subjects’ brain, and 400 whole-brain images were collected in one functional scan. The first two images of each scan were excluded from the analysis because of non-steady state magnetization. Ventricles and white matter signal were sampled using a grow-region function embedded in the Brain Voyager from a seed in each individual brain. Using MATLAB (MathWorks, Natick, MA) ventricle and white matter time-courses were regressed out of the data and the resulting time course was filtered to the frequency band-width of 0.1-0.01 Hz (in which typical spontaneous BOLD fluctuations occur). The resulting data were then imported back onto Brain Voyager for further analyses. Single subject data were registered to cortical space (as was done with the task data), and were spatially smoothed with a two-dimensional 4 vertex half-width Gaussian. The seed regions-of-interest (ROI) was defined for the sensorimotor hand region (100% overlap of controls individual activation for hand movement, p < 0.001 each). Individual time courses from this seed ROI were sampled from each of the participants, z-normalized and used as individual predictors in single-subject GLM analyses. For each group, functional connectivity parameter estimate values were sampled from regions showing full overlap probability of individual subjects’ activation for each other body part (right foot, abdomen right shoulder and mouth), and averaged by group (**Fig. 2G**). for group comparison of FC patterns, ANOVA, t-test and Bayesian statistics were used as in for task activation comparisons. The Bayes factor (BF_10_) was calculated from computing a two-samples independent t-test between the groups on FC to the hand area (**Fig. 3B**). Additionally, functional connectivity between the right and left sensorimotor hand ROIs and correlation to the global signal (e.g. (2)) were compared between the groups. The right hemisphere hand ROI defined by 100% subjects’ activation for moving the left hand in the controls, comparable to the definition of the left hemisphere ROI. Bilateral functional connectivity between the hand ROIs did not differ between the groups (p > 0.55), in accordance with the bilateral sensorimotor deprivation. Functional connectivity to the global signal was assessed by correlating the time-course of the hand ROI with the averaged resting-state time-course across all grey-matter voxels, at a single subject level. No difference in correlation was found between the groups (p > 0.10), although the relatively low p value suggests the group size may not permit conclusive findings in this case. No group difference was found also when computing correlation with the global signal excluding the sensorimotor strip (p > 0.49).

## Author Contributions

E.S.-A., G.V., and A.C. designed research; E.S.-A. and G.V. performed research; E.S.-A. analyzed data; and E.S.-A., G.V., and A.C. wrote the paper.

## Acknowledgements

We are thankful to the dysplasic subjects who participated in our experiments. We thank Himanshu Bhat and Thomas Benner of Siemens Healthcare for the SMS-EPI sequence, and Steven Cauley of Massachusetts General Hospital for modifications that enabled implementation of our protocols in a routine session. We thank Tamar Makin for her helpful comments on an earlier version of the manuscript. This work was supported by Società Scienze Mente Cervello–Fondazione Cassa di Risparmio di Trento e Rovereto, by a grant from the Provincia Autonoma di Trento, and by a Harvard Provostial postdoctoral fund (to A.C.); and by the European Union’s Horizon 2020 Research and Innovation Programme under Marie Sklodowska-Curie Grant Agreement 654837 and the Israel National Postdoctoral Award Program for Advancing Women in Science (to E.S.-A.).

## Supplemental Information

**Figure S1:**
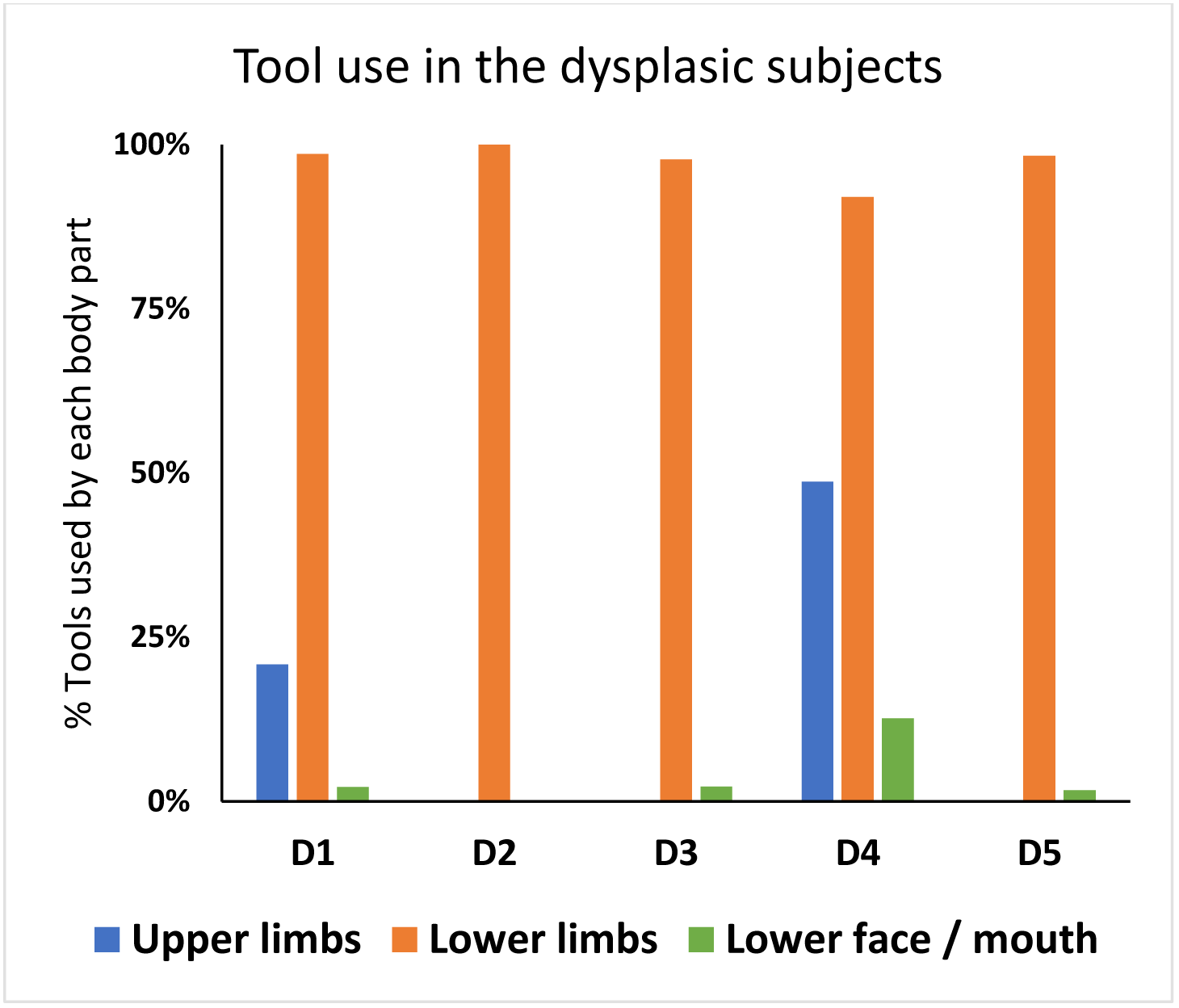
Everyday tool use by the dysplasic subjects, Related to Figure 1. The dysplasics were provided, over a month before the scan, with a list of 187 tools and small graspable objects and noted for each which body part they use if with (upper limbs, lower limbs, mouth, multiple items can be marked for the same item) or if they have never used it before to achieve its typical function. The figure depicts the percentage of using each body part, for the items they reported to have used before, for each dysplasic individual. All the dysplasic subjects reported to use tools with their lower limbs for the clear majority of tools they have experienced using. Foot tool-use accounted for a minimum of 92% of the used tools, although some tools were jointly manipulated by the lower face or remaining upper limbs in specific individuals (in subjects D1 and D4).

**Figure S2:**
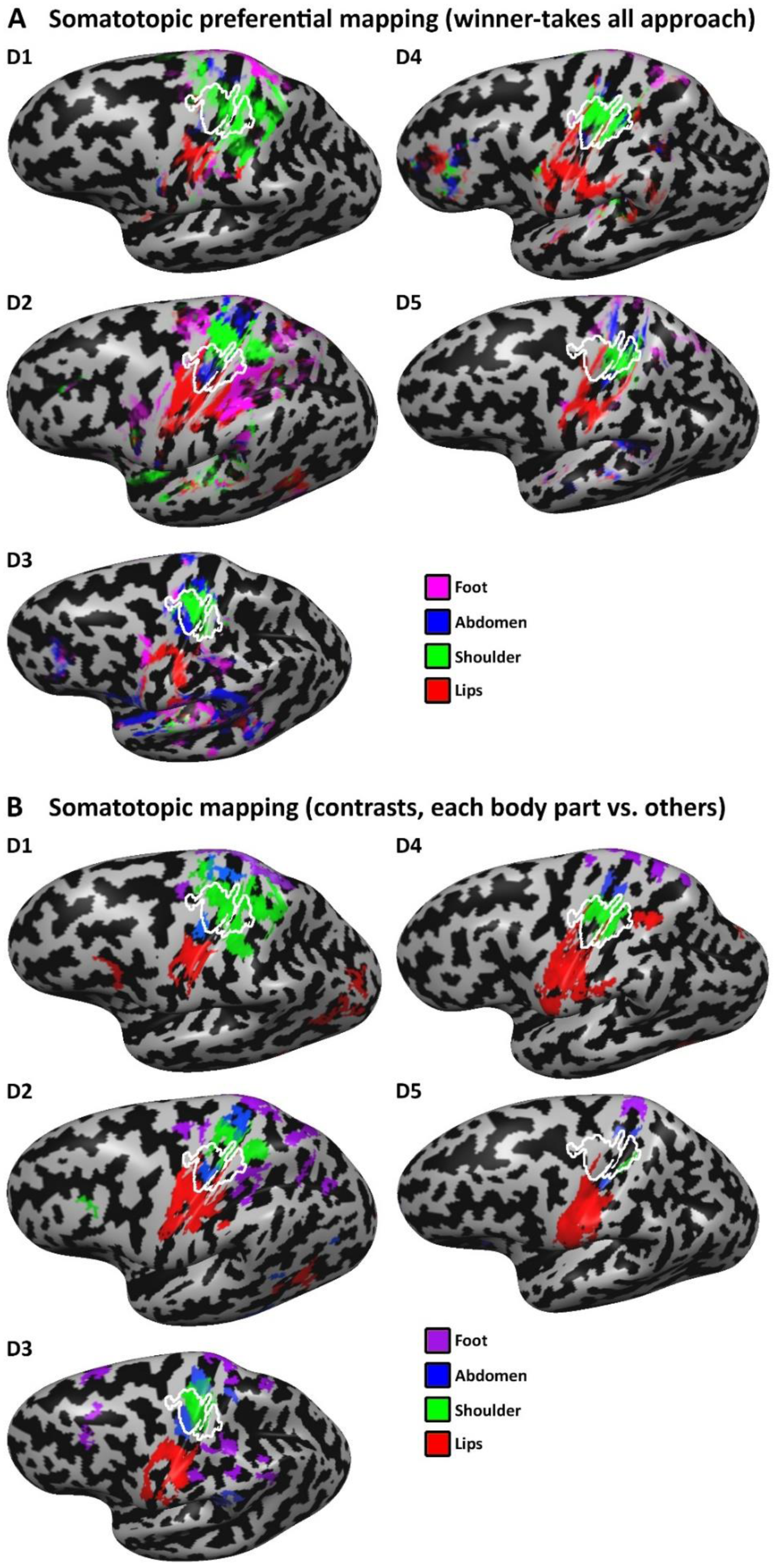
Somatotopic mapping in individual dysplasic subjects, Related to Figure 2. Preferred body part responses for flexing movements for the dysplasic individuals (unsmoothed data) largely replicates the group patterns (**Fig. 2B**), showing a preference of the lateral sensorimotor cortex (the hand area is delineated in white) to movements of the shoulder and abdomen and not of the foot, despite the extensive use of the feet to perform typically manual fine-motor tasks. Findings are presented for a winner-takes-all analysis (each vertex is colored according to highest activation; **A**) and in GLM contrasts of each body part vs. the remaining body parts (**B**; e.g. green represents a significant contrast of shoulder > foot, abdomen and lips; p < 0.05 corrected) in the individual unsmoothed data. The sensorimotor hand area, delineated in white, represents the area activated by right hand movement in all (100%) of the control participants, aligned to the dysplasics cortices according to the pattern of cortical folding.

**Figure S3:**
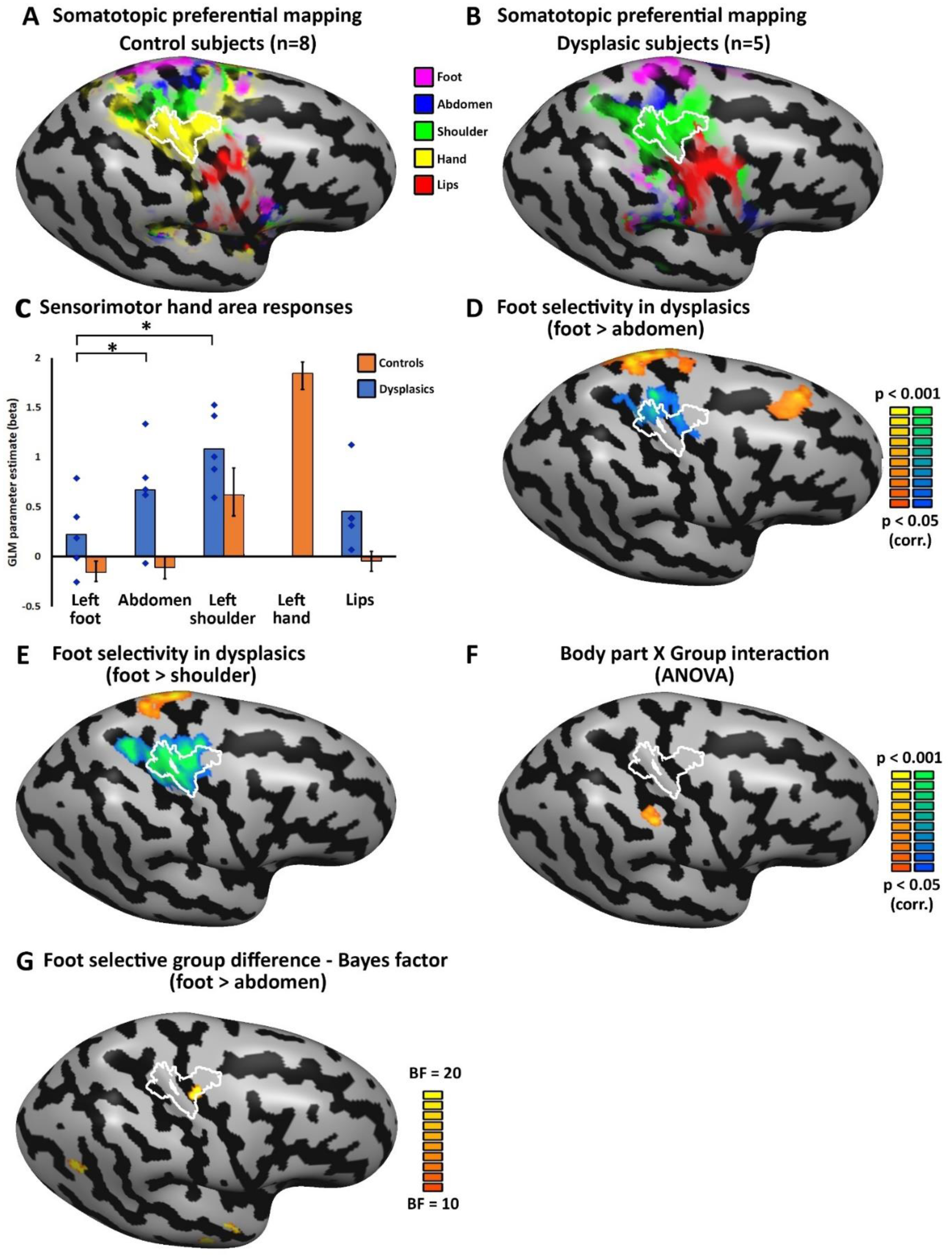
The sensorimotor hand area in the dysplasics does not show selectivity for the foot, right hemisphere (RH) data, Related to Figure 2. **A.** Preferred body part responses for contraction movements (winner-takes-all approach) for the control subjects follows the standard Penfield homunculus. The RH sensorimotor hand area is delineated in white, representing the core area activated by left hand movement in all (100%) of the control participants (each at p < 0.05 corrected), to account for inter-subject variability. **B.** Preferred body part responses for flexing movements for the dysplasic group shows a preference for shoulder movements in the RH hand area, replicating the findings in the left hemisphere, despite the use of the right foot compensatorily as an effector. **C.** Sensorimotor responses were sampled from the RH hand area, showing that this region in the dysplasics is more activated by proximal body parts (shoulder; p < 0.005 and abdomen/trunk; p < 0.01) than by foot movements. Error bars for the control group (orange bars) represent standard error of the mean. Individual data points (blue diamonds) are presented for the five dysplasic individuals in addition to the group average. **D.** Foot movement selectivity (over abdomen movement in the dysplasics can be found in the superior frontal cortex, but not in the hand primary sensorimotor cortex, which shows the reverse preference. **E.** Movement selectivity comparing the shoulder and foot in the dysplasics shows a robust preference to shoulder movement (a proximal, non-compensatory body part) rather than to foot movement in the hand area, replicating the findings in the left hemisphere. **F.** Overall body part selectivity (comparing movement of all shared body parts; e.g. lips, shoulder, abdomen and foot) differs between the dysplasics and controls (ANOVA Body part X Group interaction) in the inferior parietal lobule. **G.** Bayes factor (BF_10_) for difference between the groups in their differential activation to left foot movement (vs. abdomen movement) is shown. The dysplasics show different selectivity level for left foot movement as compare to the controls in three cortical loci, including the sensorimotor hand area. However, the group difference in found in the primary sensorimotor hand area in this analysis reflects a preference of the dysplasics group towards the abdomen movement (compare to panel **D**). A direct comparison of the selectivity to left foot movement (vs. abdomen movement) between the dysplasics and control subjects using frequentist analysis did not yield significant results.

**Figure S4:**
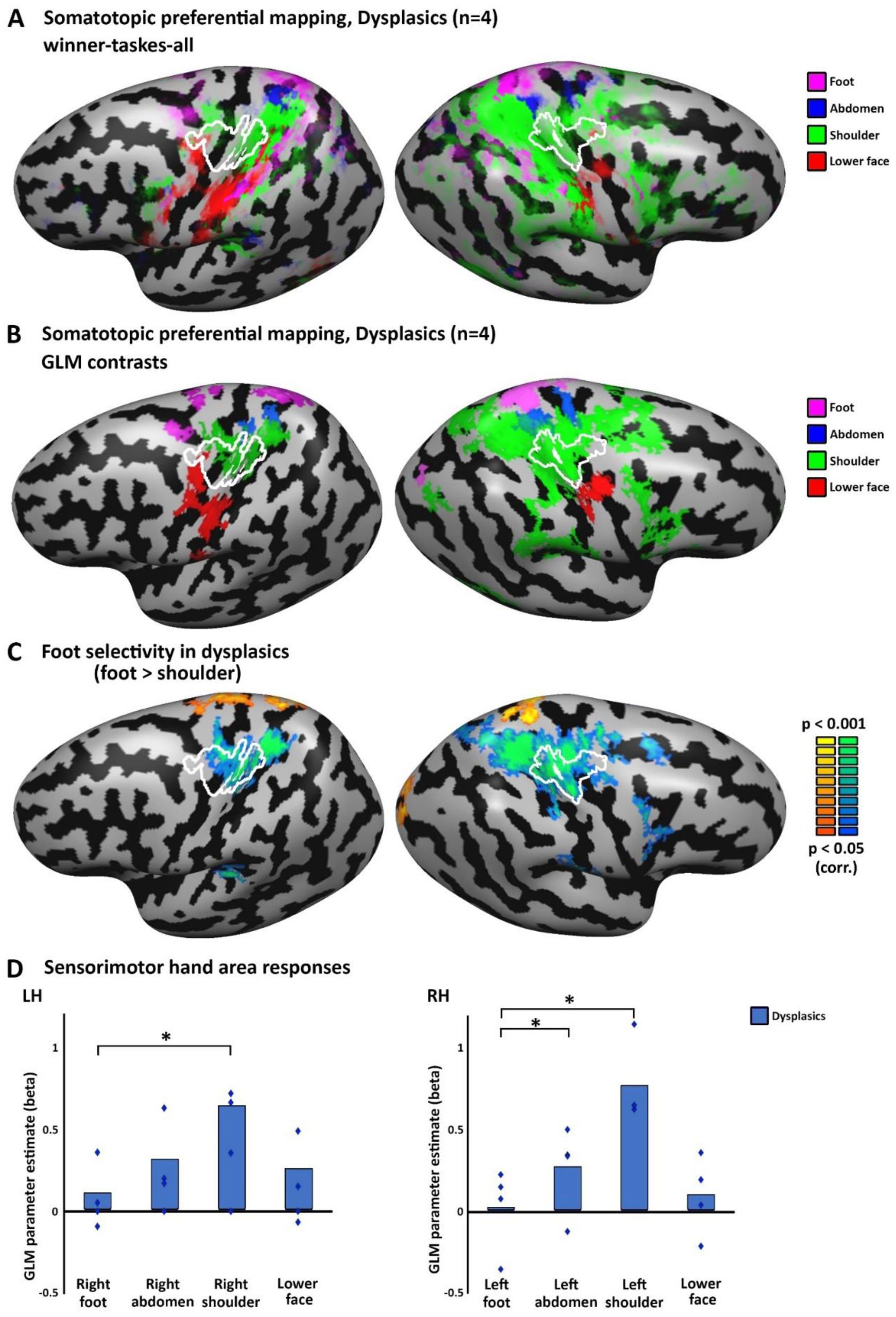
Selectivity for the shoulder in the hand area of the dysplasics in passive tactile stimulation, Related to Figure 2. **A-B.** Preferred body part responses for tactile stimulation for the dysplasic group shows a preference for contralateral shoulder movements in the hand area in both hemispheres, despite the extensive use of the feet (mainly right foot) to perform typically manual fine-motor tasks. Findings are presented for a winner-takes-all analysis (each vertex is colored according to highest activation in the group average; **A**) and in GLM contrasts of each body part vs. the remaining body parts (**B**; e.g. green represents a significant contrast of shoulder > foot, abdomen and lips; p < 0.05 corrected). The sensorimotor hand area, delineated in white, represents the area activated by right hand movement in all (100%) of the control participants. **C**. The hand primary sensorimotor cortex shows robust selectivity for shoulder over foot in the dysplasics also in passive tactile stimulation. **D.** Sensory responses were sampled from the sensorimotor hand areas in both hemispheres, showing that these regions in the dysplasics are more activated by the shoulder than by foot tactile stimulation (p < 0.005 for both comparisons, left foot > left abdomen is also significant at p < 0.05). Individual data points (blue diamonds) are presented for the five dysplasic individuals in addition to the group average (blue bars).

**Figure S5:**
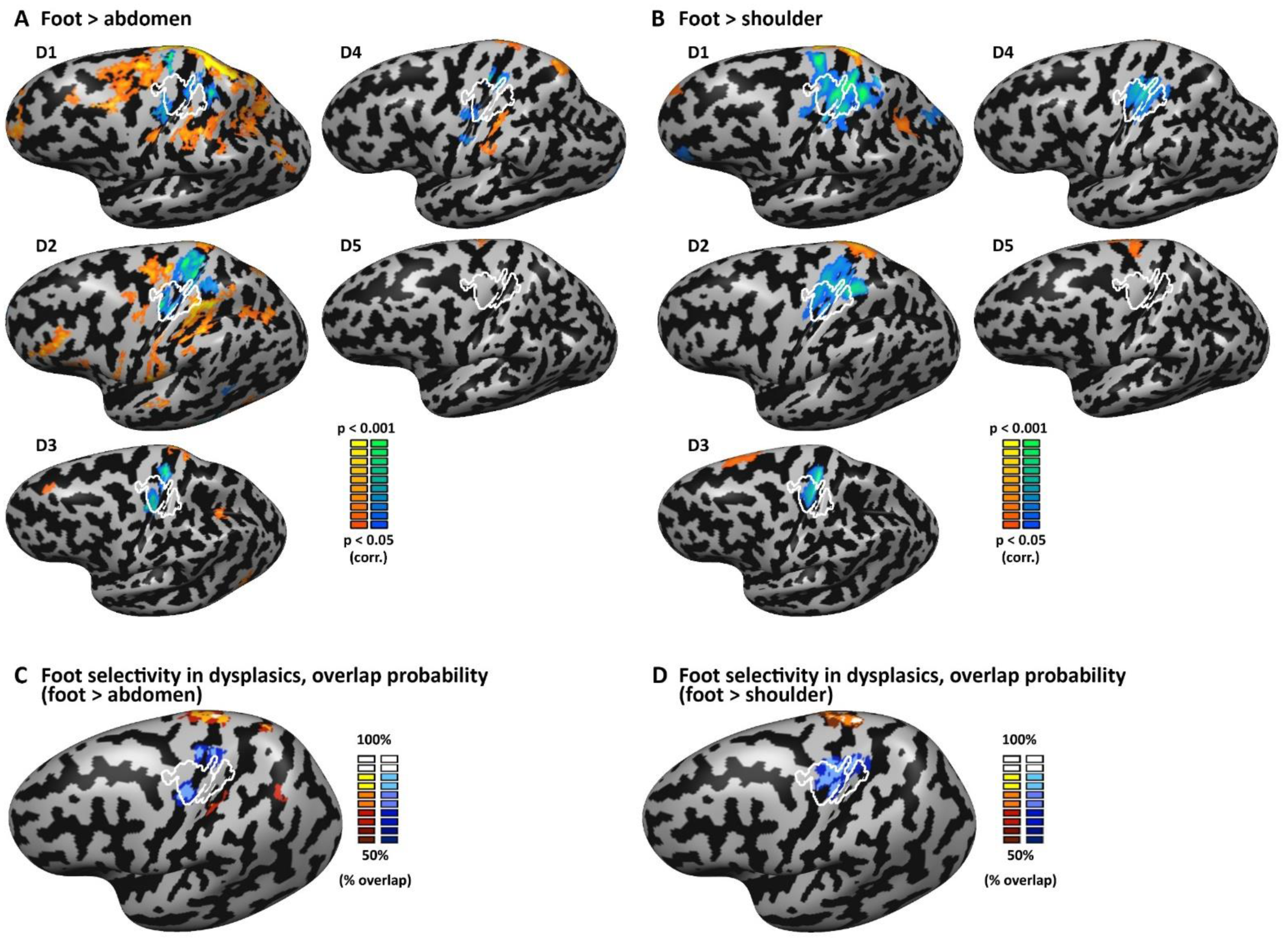
Selective foot activation in the dysplasic individuals, Related to Figure 2. **A.** Foot movement selectivity (over abdomen movement) in the dysplasic individuals (unsmoothed data, p < 0.05 corrected) can be found among other areas in the superior parietal lobule and premotor cortex, but almost nowhere (apart from small cluster in subject D2) in the hand primary sensorimotor cortex (delineated in white). In contrast, most subjects show significantly higher activity for abdomen movement in the hand region. The sensorimotor hand area, delineated in white, represents the area activated by right hand movement in all (100%) of the control participants, aligned to the dysplasics cortices according to the pattern of cortical folding. **B.** Even more robustly, the hand sensorimotor cortex of nearly all individual dysplasics shows significant preference for shoulder movement over foot movement. **C.** Probabilistic mapping of the individual subject activation for foot movement over abdomen movement (calculated from the maps appearing in panel **A**) further attests to the consistency of the preference for abdomen movement in the hand area. Each cortical vertex is colored according to the percentage of subjects showing significant activation (red) or deactivation (blue). **D.** Similarly to C, Probabilistic mapping of the individual subject activation for foot movement over shoulder movement (calculated from the maps appearing in panel **B**) further attests to the consistency of the preference for shoulder movement in the hand area.

**Table S1:** List of tools all five dysplasic subjects reported to have already used to achieve their typical function with their lower limbs, and with them only*, Related to Figure 1

|  |  |  |  |  |  |
| --- | --- | --- | --- | --- | --- |
| Bowl scraper | Cooking strainer | Glue stick | Kitchen sponge | Razor | Syringe |
| Calculator | Correction pen | Hair brush | Match | Rolling pin | Tambourine |
| Can opener | Elastic band | Hairdryer | Nail | Scissors | Thermometer |
| Cards | Erasing gum | Hand fan | Nail polish | Screw | Toaster |
| Chess pawn | File | Hole punch | Paper clip | Sewing needle | Toothbrush |
| Comb | Frisbee | Iron | Pencil sharpener | Spinning top | Vegetable peeler |
| Computer mouse | Garlic press | Kettle | Protractor | Stapler | Yo-yo |

\*The instruction the dysplasic subjects were given was the following:
Please indicate your experience in using the listed objects by putting X in the appropriate column (columns were “I use it with my upper limb(s)”; “I use it with my lower limb(s)”; “I use it with my mouth”; “I have never used it to achieve its typical function”; “I would be able to use it to achieve its typical function, if I had the opportunity to try”; “?”). If you have already used the object to achieve its typical function (e.g., using a hammer to put on a nail, using a sword to sword fight and so on), please indicate whether you used your upper limbs, lower limbs or mouth to use it. If you use an object with a combination of several body parts or if you use indifferently different body parts to use it, please put X in all the appropriate columns (for instance lower limbs and mouth). If you have never used a given tool to achieve its typical function, that is, if you have never touched it or if you have only transported it, then put X in the column “I have never used it to achieve its typical function” and, then, indicate whether you estimate that you would nevertheless be able to use it to achieve its typical function if you were given the opportunity by putting a X in the last column, or not, by letting the last column empty. If you don’t know the object, or if you are not sure of what it refers to, or if you are not sure of your response, put a X in the column “?”.

